# Evolutionary optimization of allosteric activation by Cl^-^ and Cl^-^ conduction in vesicular glutamate transporters

**DOI:** 10.64898/2026.02.25.707697

**Authors:** Victor Lugo, Yannick Güthoff, Suryapriya Ulaganathan, Arne Franzen, Sabine Balfanz, Arnd Baumann, Ghanim Ullah, Christoph Fahlke

## Abstract

Vesicular glutamate transporters harness proton gradients to load glutamate into synaptic vesicles, while mediating luminal chloride efflux through a channel-like conductance. We combined heterologous expression in mammalian cells, whole-cell patch-clamp recordings and mathematical modeling to functionally characterize the *Drosophila melanogaster* vesicular glutamate transporter DVGLUT and to compare it to a rat counterpart, rVGLUT1. As in mammalian VGLUTs, DVGLUT glutamate transport is coupled to proton exchange, in a 1:1 stoichiometry. In both, luminal Cl^-^ is necessary as allosteric activator, however, the allosteric affinity is higher in fly than in rat transporters. DVGLUT anion channels exhibit lower unitary currents, but higher anion channel open probabilities, resulting in larger Cl^-^ currents for the fly transporter in the presence of cytoplasmic glutamate. The higher allosteric affinity together with the enhanced anion channel activity may serve as evolutionary adaptation of VGLUTs to lower ion concentrations in *Drosophila*, illustrating the impact of these particular features for synaptic vesicle filling.

## INTRODUCTION

Glutamate is the major excitatory neurotransmitter in the mammalian central nervous system. Its exocytotic release by presynaptic nerve terminals relies on vesicular glutamate transporters (VGLUTs) that fill synaptic vesicles with glutamate^1–4^. There exist three mammalian VGLUT isoforms, VGLUT1, VGLUT2 and VGLUT3, with different localization and slight variation in function^5–9^. VGLUTs transport glutamate in a carrier-mediated manner, harnessing the transmembrane H^+^ gradient across the vesicular membrane for effective glutamate accumulation in synaptic vesicles. They are dual function transport proteins that can also assume anion channel modes^10–13^.

Vesicular glutamate transporters also exist in non-mammalian species, and the comparison of such transporters with mammalian counterparts promise insights into the evolutionary optimization of functional transporter features. The fruit fly *Drosophila melanogaster* has been a model system for the study of neuronal function as well as neurological dysfunctions for many decades^14,15^. with the *Drosophila* neuromuscular junction greatly contributing to our knowledge of the function, the plasticity and the development of glutamatergic synapses^16–19^. *Drosophila melanogaster* contains only a single vesicular glutamate transporter, DVGLUT, that is expressed in synaptic vesicles of glutamatergic motoneurons and interneurons and in synaptic terminals of glutamatergic neuromuscular junctions (NMJs)^20^.

We here used heterologous expression of a variant of DVGLUT with removed intracellular retention signals^11,12,21^ and whole-cell patch clamp recordings for a detailed functional analysis of the fly vesicular glutamate transporter. The comparison of transport functions of the two orthologues demonstrated that VGLUT glutamate transport is conserved between the two species, whereas VGLUT activation by luminal anions and anion channel function differs between fly and rat in an ortholog-specifc manner.

## Results

### DVGLUT functions as voltage-, pH- and chloride-dependent anion channels in the absence of glutamate

Removing intracellular retention signals results in effective surface membrane insertion of rat vesicular glutamate transporters in *Xenopus* oocytes and mammalian cells^11,12^. The homologous amino acid exchanges (Y38A/E39,40AA/M41A/E42/G43,44AA) promoted surface expression of DVGLUT (DVGLUT_PM_, Figure S1) after heterologous expression in mammalian cells and made functional analysis by whole-cell patch clamp possible also for the fly vesicular glutamate transporter. Since VGLUTs require luminal acidification to be active, their functional characterization is complicated by proton-activated anion (PAC) channels that are endogenously expressed in most cell lines^22,23^. PAC channels are activated by positive membrane potentials under acidic pH conditions and thereby restrict the voltage range, over which VGLUT currents can be reliably separated from background conductances. We generated a mutant HEK293T cell line (KO_PAC_HEK293T) , in which these proton-activated anion (PAC) channels were knocked-out using the CRISPR-Cas9 system (Figure S2) and used this cell line for the majority of experiments.

In the absence of the transport substrate glutamate, VGLUTs exclusively function as anion channel. Figure 1A depicts representative Cl^-^ current recordings of a KO_PAC_HEK293T cell expressing DVGLUT_PM_. Cells were dialyzed with a CholineCl-based intracellular solution at pH_i_ 7.4, and perfused with CholineCl-based extracellular solutions of different pH_o_ values. At extracellular pH 7.4, current amplitudes were indistinguishable from untransfected cells (−32 ± 4 pA at -160 mV for DVGLUT_PM_, *n* = 32; and -39 ± 2 pA at -160 mV for untransfected cells, *n* = 10). At acidic pHs, hyperpolarizing voltage steps elicit current activation, resulting in amplitudes up to 3.1 ± 0.5 nA (at -160 mV and pH_o_=5.0 (*n* = 34)). In the following, we provide – if not stated otherwise – proton-activated current amplitudes after correction for leakage by subtracting current values obtained at the same cell at pH_o_ 7.4 or higher pH_o_ values at otherwise identical conditions. Figure 1B provides the pH and voltage dependence of normalized DVGLUT_PM_ currents.

**Fig. 1.**
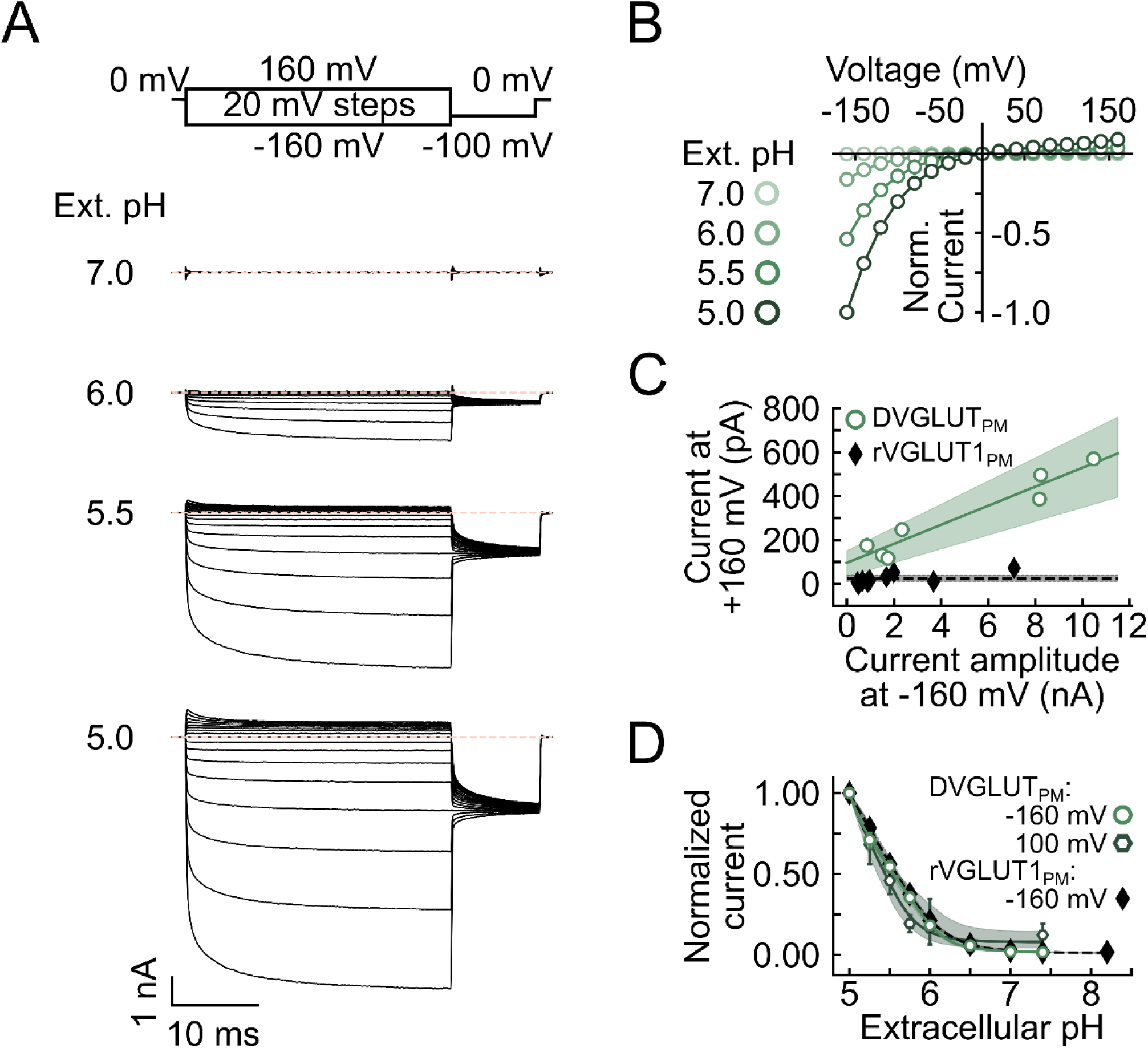
pH- and voltage-dependent gating of DVGLUT_PM_ anion channels. **(A)** Representative recordings from a KO_PAC_HEK293T cell expressing DVGLUT_PM_ at four different external pH values. The cell was dialyzed with a Cl⁻-based intracellular solution at neutral pH and perfused with extracellular solutions at the indicated pH values. The dashed line indicates the zero-current level. (**B)** Steady-state voltage dependence of background-subtracted currents after normalization to the current amplitude at −160 mV (pH_o_ 5.0, *n* = 32; pH_o_ 5.5, *n* = 29; pH_o_ 6.0, *n* = 25; pH_o_ 7.0, *n* = 25). **(C)** Plots of current amplitudes at +160 mV versus current amplitudes at −160 mV for individual cells expressing DVGLUT_PM_ (open symbols) or rVGLUT1_PM_ (filled symbols). Lines and shaded areas indicate the mean and 95% confidence interval (CI) derived from a sloped linear fit (DVGLUT_PM_, *r*² = 0.93) or a zero-slope fit (rVGLUT1_PM_). (**D)** Dose–response plots of steady-state current amplitudes measured at −160 mV (filled symbols, rVGLUT1_PM_; open symbols, DVGLUT_PM_) and at +100 mV (open symbols with dark outlines, DVGLUT_PM_). Lines indicate the mean fits obtained with hyperbolic-type models; shaded areas represent the 95% confidence interval (CI).

At positive voltages, DVGLUT_PM_ currents are small, with time-dependent deactivation (Figure 1A). Figure 1C depicts plots of current amplitudes at +160 mV versus valuess at -160 mV for multiple cells expressing either DVGLUT_PM_ (open symbols) or rVGLUT1_PM_ (filled symbols). These data can be fit with a linear relationship with positive slope for DVGLUT_PM_ (solid line, *r*^2^ =0.93), whereas the slope is not different from 0 for rVGLUT1_PM_ (dashed lines, p **=** 0.015, F-test results with alpha = 0.01). Figure 1D provides the pH dependence of DVGLUT_PM_ Cl^-^ currents at -160 mV and at +100 mV, with closely similar dose-response curves for negative and positive voltages. At negative voltages, pH dependences of rat and fly VGLUT anion channels resemble each other (Figure 1D).

Whereas rVGLUT1_PM_ is closed at 0 mV and at positive voltages^12^, DVGLUT_PM_ deactivates to current amplitudes clearly above background at positive potentials (Figure 1B). Figure 2A depicts representative current responses of DVGLUT_PM_ (top) and rVGLUT1_PM_ (bottom) to two step protocols with an initial voltage step from 0 mV to -160 mV to activate VGLUT anion channels. During this step, the fly channel (black) opens faster than its mammalian counterpart (gray) (Figure 2B). Subsequent steps to positive voltages elicit current decay in both fly and mammalian transporters (Figure 2C). Both, closing time constants (Figure 2D) and the remaining residual outward currents (Figure 2E) are significantly larger for DVGLUT_PM_ (green) than for VGLUT1_PM_ (gray).

**Fig. 2.**
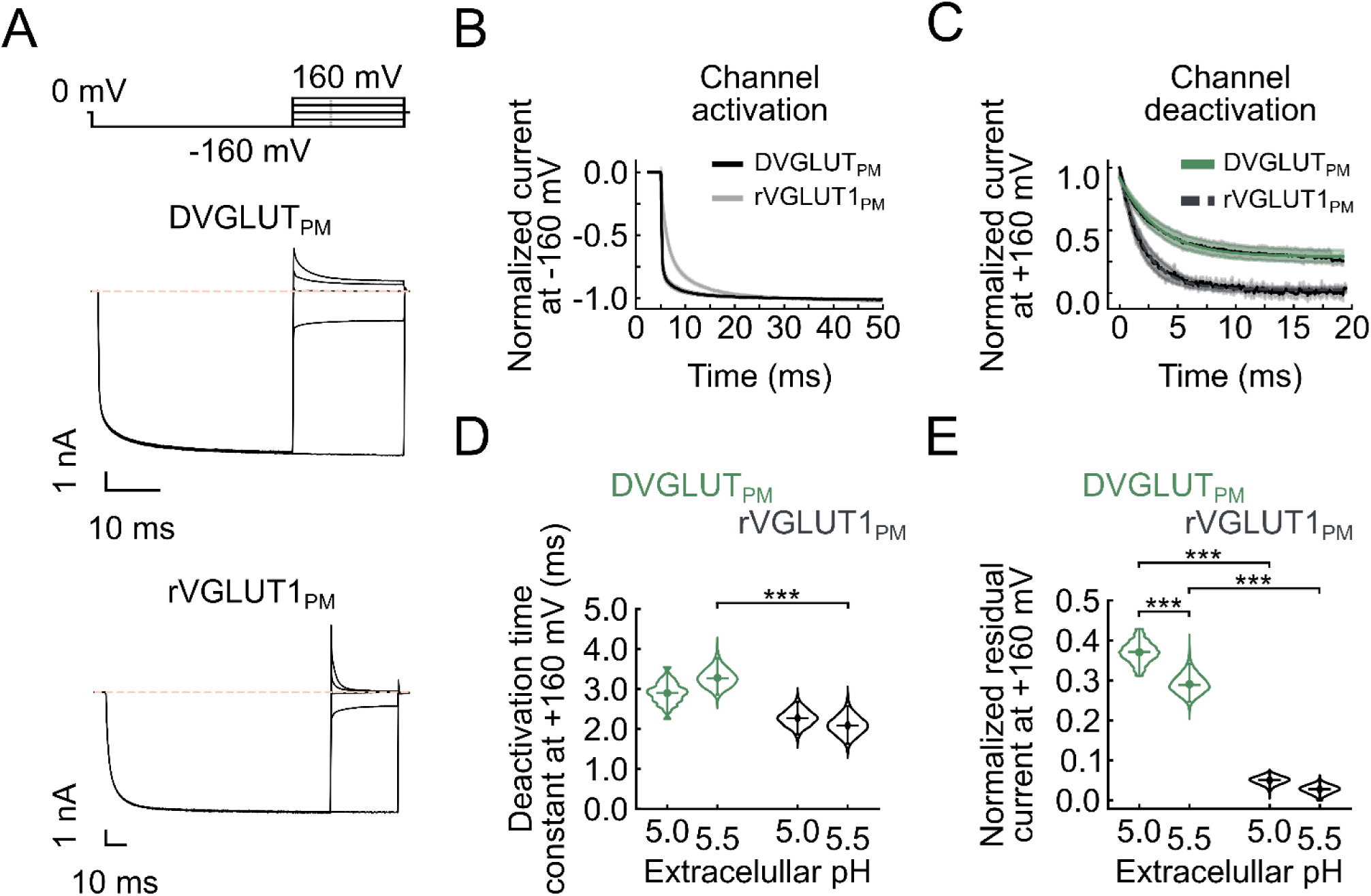
Fly and rat VGLUT_PM_ anion channels differ in voltage-dependent gating. **(A)** Representative current responses of KO_PAC_HEK293T cells expressing DVGLUT_PM_ (top) or rVGLUT1_PM_ (bottom) to a pulse protocol consisting of a fixed −160 mV prepulse followed by steps to various voltages. Cells were dialyzed with a Cl⁻-based intracellular solution at neutral pH, and currents were recorded at pH_o_ = 5.0. The dashed line indicates the zero-current level. (**B, C)** Mean values and 95% confidence intervals (CI) of the activation (**B**) or deactivation (**C**) time courses for normalized DVGLUT_PM_ (*n* = 7) and rVGLUT1_PM_ (*n* = 22) chloride currents, recorded in experiments as shown in **A**. (**D, E)** Time constants (**D**) and normalized residual currents (**E**) obtained from monoexponential fits to recordings as shown in panel C at pH_o_ 5.0 (DVGLUT_PM_, *n* = 4; rVGLUT1_PM_, *n* = 9) and pH_o_ 5.5 (DVGLUT_PM_, *n* = 15; rVGLUT1_PM_, *n* = 11).

Vesicular glutamate transporters are activated by luminal Cl^-21,24–26^. Figure 3A shows representative DVGLUT_PM_ whole-cell recordings upon consecutive perfusion with 0 mM, 2.2 mM and 140 mM external [Cl^-^] at pH_o_=5.0, with Figure 3B depicting normalized late chloride currents elicited at -160 mV at these chloride concentrations. Plots of the normalized maximum currents at -160 mV and pH 5.0 versus external [Cl⁻] were fit with single dose–response relationships using Hill coefficients fixed to 1, yielding a voltage-independent EC_50_ of 2.5 ± 0.4 mM (*n* = 17) (Figure S3), lower than the mammalian rVGLUT1_PM_ (EC_50_(rVGLUT1_PM_) = 34.2 ± 17.9 mM, *n* = 11, *p* < 0.01) (Figure 3C). Higher external [Cl^-^] enhances DVGLUT_PM_ currents, but fly as well as rat VGLUT_PM_ anion channels are also active in the absence of external Cl^-^ ^13^. We therefore fitted a Cl^-^-independent component to account for this property. Chang et al.^21^ recently identified a highly conserved arginine in the outer vestibule (R176) as main determinant of the allosteric Cl^-^ activation of rVGLUT1. Neutralization of the corresponding residue (R199) in DVGLUT_PM_ increases the Cl^-^-independent component (from 0.33 ± 0.02 in WT, n = 17; to 0.76 ± 0.04 in R199A DVGLUT_PM_, n=17, *p* < 0.001), but leaves its apparent affinity for luminal chloride unaltered (R199A, EC_50_= 6.4 ± 3.7 mM, p > 0.1, Figure 3C).

### DVGLUT anion channels exhibit a lyotropic anion selectivity

Rat VGLUT1 is permeable to various anions with lyotropic anion selectivity, preferring large and polyatomic anions^12^. Figure 4 compares representative DVGLUT_PM_ recordings from cells with Cl^-^, NO_3_ ^-^ and I^-^ as intracellular anions. Changes in permeant anions affect the time and voltage dependence of currents. Upon hyperpolarizing voltage steps, DVGLUT_PM_ NO ^-^ currents activate with a faster time course than Cl^-^ currents; I^-^ currents increase instantaneously in response to hyperpolarizing voltage steps, followed by a slight time-dependent decrease in current amplitudes (Figure 4A). Figure 4B provides plots of steady-state current amplitudes at -160 mV of individual DVGLUT_PM_ transfected cells dialyzed with Cl^-^, NO_3_ ^-^ or I^-^-based intracellular solutions versus whole-cell fluorescences. DVGLUT_PM_ was expressed as eGFP fusion protein, and whole-cell fluorescences thus provide a measure for expression levels of individual cells. We always measured fluorescences before establishing the whole-cell configuration, to prevent possible modifications of the subcellular transporter distribution by the intracellular anion^27,28^. The linear relationship between whole-cell current amplitudes and expression levels demonstrated that intracellular trafficking did not saturate in any of the cells; the number of transporters in the surface membrane is therefore proportional to the total number of expressed transporters. Whole-cell fluorescences thus provide a relative measure of the number of DVGLUT_PM_ in the surface membrane, and the slopes of linear fits give relative current amplitudes corrected for differences in DVGLUT_PM_ expression between separate cells. The voltage dependence of DVGLUT_PM_ current amplitudes normalized to whole-cell fluorescence of the same cell reveal an anion conductance sequence of NO₃⁻ > I⁻ > Cl⁻ (Figure 4C). In cells expressing DVGLUT_PM_ and perfused with 40 mM extracellular chloride, currents recorded in 140 mM intracellular chloride reversed at +39 ± 3 mV. Substitution with cytosolic iodide shifted the reversal potential slightly to +43 ± 1 mV (p = 0.14), whereas replacement with nitrate produced a significant increase to +52 ± 1 mV (p < 0.001). These reversal potentials correspond to relative permeabilities (P_x_/P_Cl_) of 1.2 ± 0.1 for I⁻_i_ and 1.8 ± 0.2 for NO₃⁻_i_ (p < 0.001). As already observed for rVGLUT1 ^12^, the pH dependence of DVGLUT inward current reponses is shifted to the right by intracellular NO ^-^ and I^-^ (Figure 4D).

**Fig. 3.**
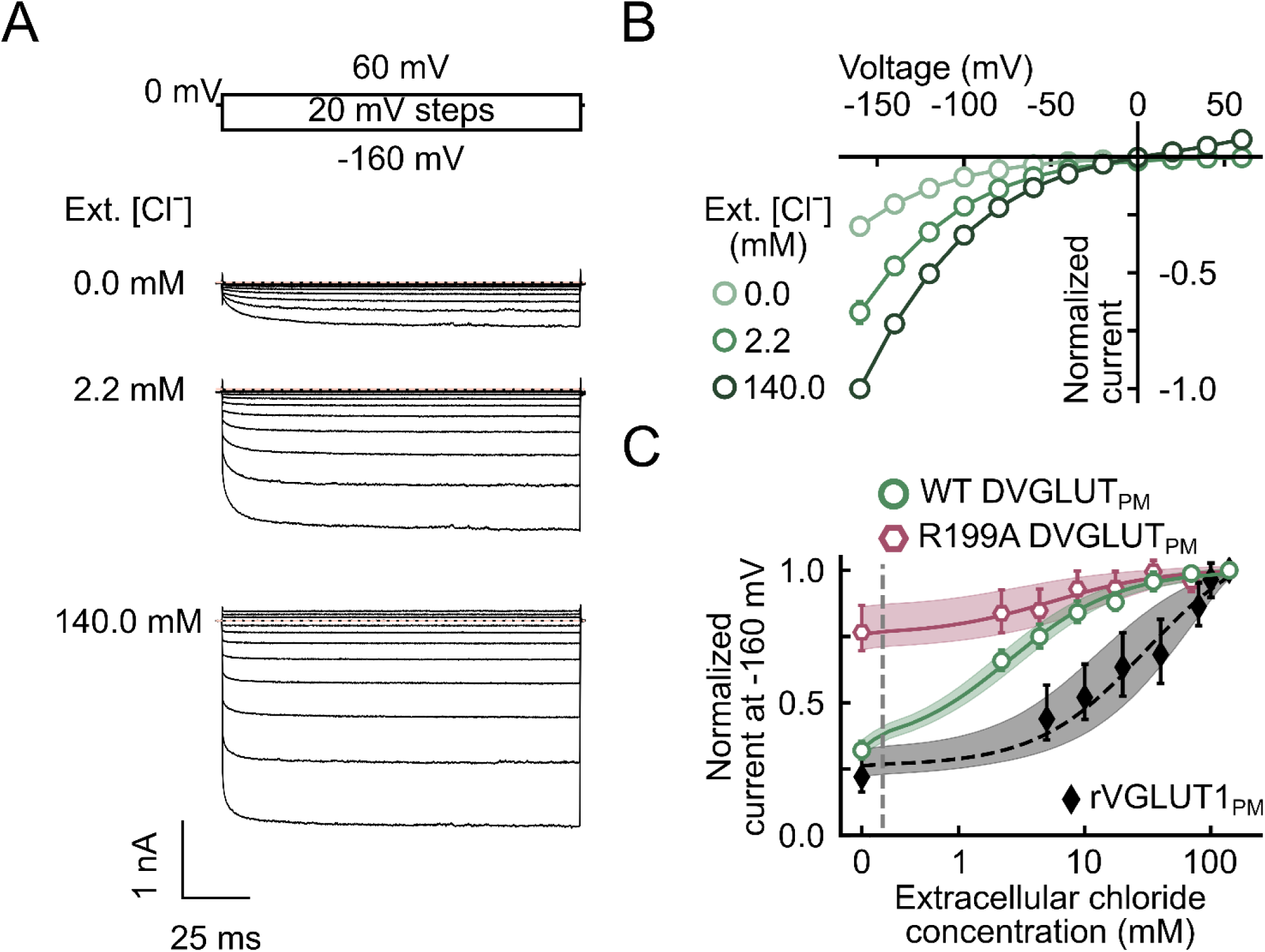
Distinct allosteric Cl⁻ dependences of DVGLUT_PM_ and rVGLUT1_PM_. **(A)** Representative recordings from Flp-In T-REx 293 cells expressing DVGLUT_PM_ at three different external Cl⁻ concentrations at pH_o_ 5.0. Cells were dialyzed with a Cl⁻-based internal solution at neutral pH and perfused with extracellular solutions containing the indicated Cl⁻ concentrations. Currents were leak-corrected by subtracting the currents recorded at pH_o_ 7.4 and [Cl⁻]_o_ = 140 mM under the same voltage protocol. Dashed line indicate the zero-current level. (**B)** Steady-state current–voltage relationships at pH_o_ 5.0 and three different extracellular [Cl⁻]_o,_ after normalization to the maximum current amplitude measured at −160 mV and [Cl⁻]_o_ = 140 mM (n = 14; pH_i_ = 7.5). (**C)** Current amplitudes at −160 mV as a function of extracellular [Cl⁻] (open circles, DVGLUT_PM_, *n* = 17; open hexagons, R199A DVGLUT_PM_, *n* = 17; filled symbols, rVGLUT1_PM_, *n* = 11). A hybrid linear–logarithmic x-axis was used; the dashed vertical line indicates the transition between linear and logarithmic scaling (0.15 mM), such that responses at 0 chloride concentration are displayed on a linear scale. No background subtraction was performed. Lines indicate the mean fits with dose–response relationships (Hill coefficient was fixed to 1); shaded areas show the 95% CI.

**Fig. 4.**
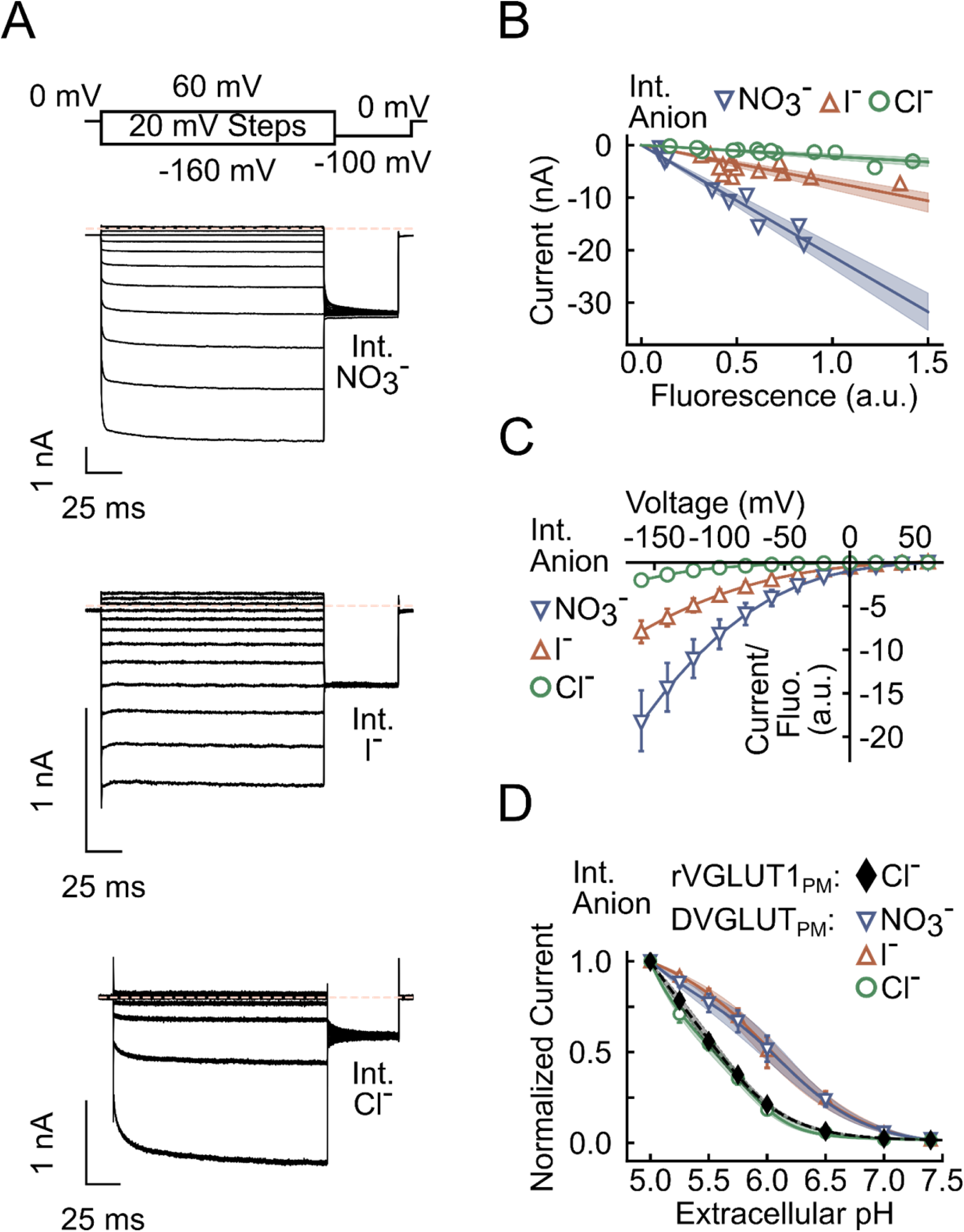
DVGLUT_PM_ anion channels exhibit a lyotropic anion selectivity sequence. ***(A)*** Representative recordings from HEK293T cells dialyzed with intracellular solutions containing Cl⁻, NO₃⁻, or I⁻ at neutral pH_i_ during voltage steps between −160 mV and +60 mV at extracellular pH_o_ 5.5. Proton-activated currents were corrected by subtracting currents values measured at pH_o_ 7.4. The dashed line indicates the zero-current level. (***B)*** Whole-cell current amplitudes measured 100 ms after a voltage step to −160 mV increase linearly with whole-cell fluorescence for cells expressing DVGLUT_PM_ and dialyzed with Cl⁻, NO₃⁻, or I⁻. The extracellular solution contained 40 mM Cl⁻ at pH_o_ 5.5. Each symbol represents background-subtracted current amplitudes from an individual cell. Lines and shaded areas indicate the mean and 95% CI from linear fits. (***C)*** Voltage dependence of background-subtracted current amplitudes measured 100 ms after the voltage step and normalized to whole-cell fluorescence values (Cl⁻, *n* = 14, NO_3_⁻, *n* = 12, I⁻ , *n* = 15). (***D)*** Dose–response plots of background-subtracted current amplitudes measured at −160 mV (filled symbols, rVGLUT1_PM_; open circles, Cl⁻; open downward triangles, NO₃⁻; open upward triangles, I⁻; DVGLUT_PM_). Solid lines indicate the mean fits to hyperbolic-type functions; shaded areas show the 95% CI.

### DVGLUT anion channels exhibit smaller unitary current amplitudes than rVGLUT1

DVGLUT anion currents are associated with Lorentzian current noise; such type of noise is typical for ion channels and is generated by the random opening and closing of individual ion channels ^29,30^. Figure 5A depicts an average power spectrum at -100 mV with intracellular Cl^-^ as charge carrier that was fitted with the sum of a pink-noise term (a = 1.3) plus two Lorentzian terms (dotted lines) with corner frequencies fc₁ = 1321 Hz and fc₂ = 85 Hz, predicting fast opening and closing of DVGLUT_PM_ anion channels. The comparison of experimental values with the prediction for uniporter-mediated shot noise (8.4 × 10⁻²⁹ A²/Hz) obtained from Schottky’s theorem (S(f) = 2 Iq, with I = 263 ± 31 pA; n = 12) reveals much larger noise amplitudes than expected for independent transporter turnover. Experiments with non-transfected HEK293T cells exhibit lower current variances and power spectra resembling 1/*f* noise under the same conditions^12,31^.

**Fig. 5.**
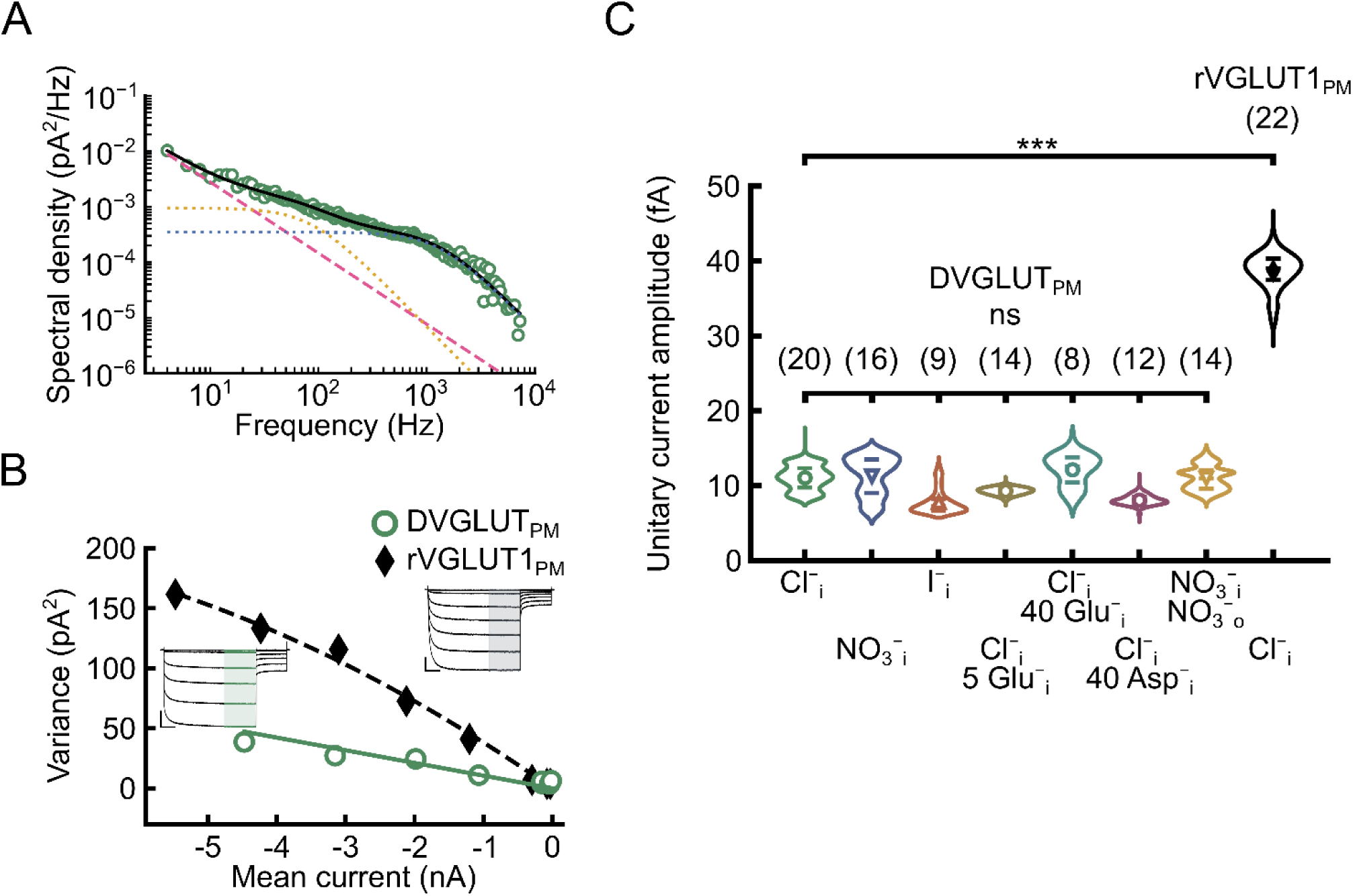
DVGLUT_PM_ anion channels exhibit smaller unitary current amplitudes than rVGLUT1_PM_. (A) Averaged power spectral density (PSD) from 12 cells expressing DVGLUT_PM_, dialyzed with Cl⁻ at neutral pH, recorded at −100 mV and pH_o_ 5.5, after subtraction of spectral densities measured at pH_o_ 7.4. The solid line represents the fit of a second-order Lorentzian function (dotted lines, S₁ = 3.5 × 10⁻²⁸ A²/Hz; S₂ = 9.5 × 10⁻²⁸ A²/Hz) in addition to pink noise (dashed line; a = 1.3). (B) Current–variance plot from representative cells expressing DVGLUT_PM_ (open symbols) or rVGLUT1_PM_ (filled symbols) and dialyzed with Cl⁻ at netural pH. The solid line represents a linear fit, the dashed line a parabolic fit. Cells were exposed to various extracellular pH values, and mean amplitudes and variances were measured at least 100 ms after voltage pulses to −160 mV (inset, scale bars: 1 nA, 25 ms). (**C)** Unitary current amplitudes of DVGLUT_PM_ or rVGLUT1_PM_ under various conditions. Number of cells in each condition are given in parentheses. In experiments with symmetrical NO_3_^-^, the external [Cl^-^] was 0.2 mM.

These results demonstrate that DVGLUT_PM_ anion current-associated noise permits the use of noise analysis to describe unitary properties of DVGLUT_PM_ anion channels. We determined current noise and current amplitudes upon variation of the external pH in order to modify open probabilities of DVGLUT_PM_ anion channels. Figure 5B depicts representative noise analysis experiments for DVGLUT_PM_ and rVGLUT1_PM_. In such experiments, cells expressing DVGLUT_PM_ or rVGLUT1_PM_ were consequently perfused with external solutions with pH values ranging from pH 8.2 to pH 5.0, and current variances were plotted versus current amplitudes and fitted with a linear (DVGLUT_PM_) or a parabolic (rVGLUT1_PM_) relationship. In nearly symmetrical chloride, such fits provide a unitary current amplitude of about 11 fA at -160 mV for DVGLUT_PM_, while its mammalian counterpart exhibits a four-fold greater amplitude (Figure 5C). Changing intracellular anions to I^-^ or NO_3_ ^-^ leaves unitary current values unmodified. Substituting 40 mM intracellular chloride with the transport substrates glutamate or aspartate (which reduces [Cl^-^]_i_ to 85 mM) did not alter DVGLUT unitary current amplitudes. Likewise, bulk substitution of extracellular chloride with 140 mM nitrate did not modify the unitary current of DVGLUT-expressing cells dialyzed with intracellular nitrate.

### DVGLUT is a coupled H^+^-glutamate exchanger

Glutamate transport is the main physiological function of the VGLUTs. Glutamate transport rates are much lower than those of Cl^-^ conduction^12^ and can therefore only be quantified after complete substitution of intracellular Cl^-^ by glutamate^-^. Figure 6A shows leakage-subtracted representative recordings from a KO_PAC_HEK293T cell expressing DVGLUT_PM_, intracellularly dialyzed with glutamate, during two consecutive perfusions at pH_o_ 5.5 and 5.0. As already reported for rVGLUT1 ^12^, intracellular dialysis with glutamate makes Cl^-^ current deactivation substantially less complete, resulting in Cl^-^ currents significantly larger than background at positive voltages. Moreover, changing the extracellular pH from 5.5 to 5.0 leads to a substantial increase of the outward Cl^-^ current at positive voltages, whereas glutamate transport currents (at negative voltages) are enhanced to a lesser degree.

**Fig. 6.**
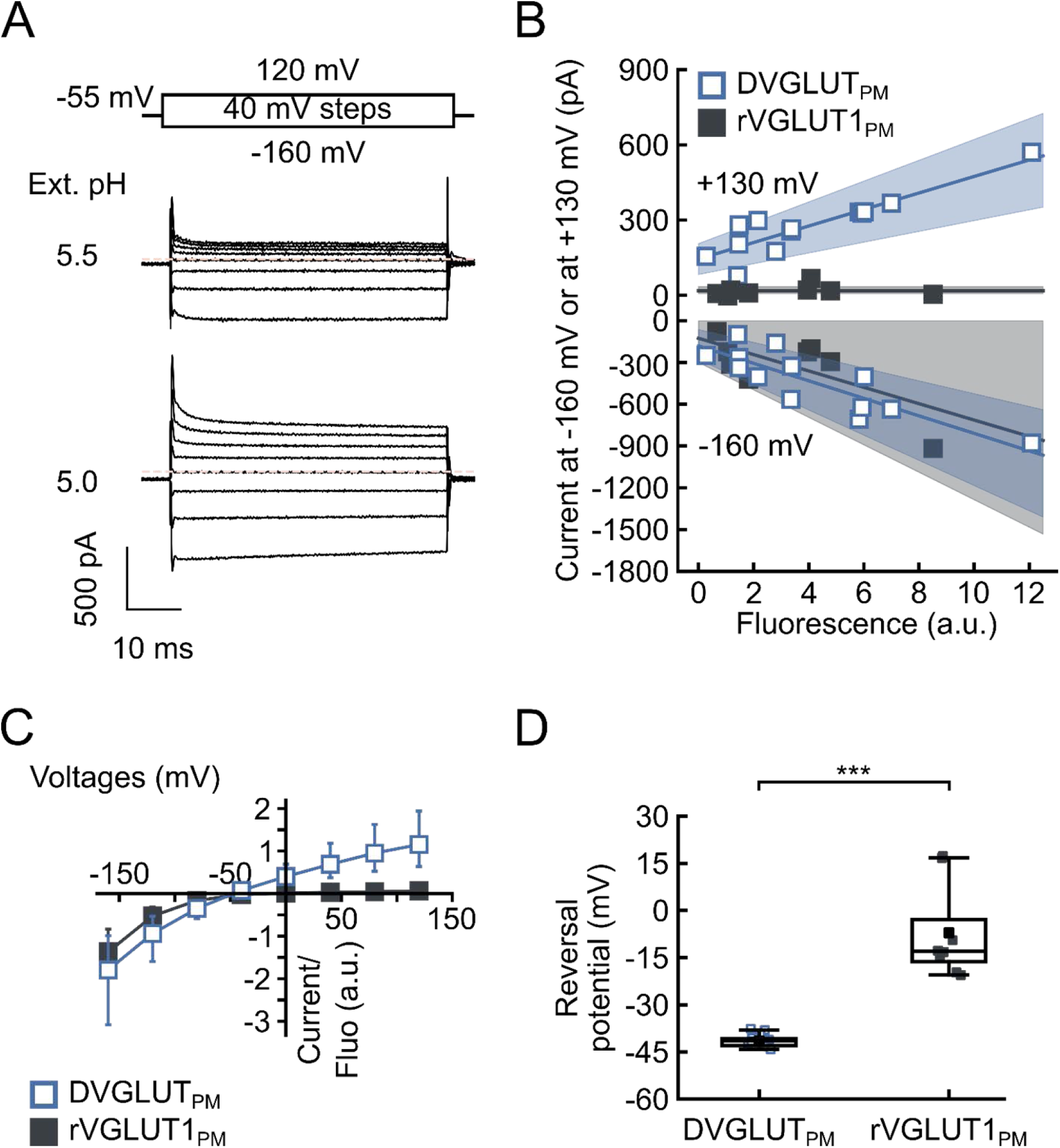
DVGLUT_PM_ anion currents under glutamate transport conditions. **(A)** Representative recordings from a cell dialyzed with 140 mM glutamate at pH 7.5 in absence of chloride or other permable anions, and perfused with 40 mM choline chloride solutions at pH 5.5 or 5.0. The dashed line indicates the zero-current level. **(B)** Plots of late whole-cell current amplitudes at −160 mV or +130 mV against whole-cell fluorescence for cells expressing DVGLUT_PM_ (open symbols) or rVGLUT1_PM_ (filled symbols). Cells were dialyzed with 140 mM glutamate and perfused with 40 mM Cl⁻ at pH_o_ 5.0. Each symbol represents current amplitudes from an individual cell. Lines and shaded areas show the mean and 95% CI from linear fits. **(C)** Current-voltage dependences for DVGLUT_PM_ (open symbols), or VGLUT1_PM_ (filled symbols). Currents were obtained from experiments as shown in **(A)**, normalized to whole-cell fluoresence intensities and presented as means ± 95% CI (DVGLUT_PM_, *n* = 13, rVGLUT1_PM_, *n* = 8). **(D)** Current reversal potentials from data shown in (C) for DVGLUT_PM;_ or rVGLUT1_PM_.

Figure 6B depicts the plot of glutamate transport currents at -160 mV and of Cl ^-^ currents at +130 mV, both at pH_o_ = 5.0, versus whole-cell fluorescences of cells, which express either DVGLUT_PM_ (open symbols) or rVGLUT1_PM_ (closed symbols). Since neither DVGLUT_PM_ nor rVGLUT1_PM_ exclusively insert into the surface membrane, the whole-cell fluorescence amplitude also includes transporters that are located in intracellular membrane compartments and thus do not contribute to the measured currents. We therefore compared fly and rat transporter distributions in confocal images of transfected cells (Figure S4). Despite some differences in the membrane distribution, with DVGLUT_PM_ exhibiting a more punctuate distribution than rVGLUT1_PM_, there are only slight differences in estimated percentages of surface membrane inserted proteins (DVGLUT_PM:_ 0.68 ± 0.01; rVGLUT1_PM:_ 0.76 ± 0.01). At comparable expression levels, carrier-mediated currents of rat and fly transporters are not different, yet DVGLUT_PM_ Cl^-^ currents are much larger than rVGLUT1_PM_ at the same fluorescence. For current-voltage relationships, we normalized current amplitudes to the whole-cell fluorescence of the same cell (Figure 6C). The larger outward Cl^-^ currents result in almost linear current-voltage relationships for DVGLUT_PM_.

Reversal potentials for glutamate currents were significantly more negative for DVGLUT_PM_ than for rVGLUT1_PM_ (Figure 6D). This difference may indicate distinct transport stoichiometries of fly and rat VGLUTs, or, alternatively, represent the consequence of the larger outward Cl^-^conductance of the fly transporter. We quantified DVGLUT_PM_ transport coupling using the same experimental approach we developed for rVGLUT1 ^12^: glutamate currents were measured at pH_o_ 5.0 and corrected for background by subtracting recordings from the same cell at pH_o_ 7.5. Current reversal potentials were measured upon variation of transmembrane proton gradients by modifying the intracellular pH between pH 6.5 and pH 8.5. To correct for the much larger Cl ^-^ inward fluxes of fly transporters, which might dominate current reversal potentials, we reduced the extracellular [Cl^-^] from 40 mM to 4 mM (Figure 7A). Measured current reversal potentials change in a linear fashion with the Nernst potential for H^+^ (Figure 7B), indicating that glutamate transport by DVGLUT is stoichiometrically coupled to H^+^ transport. The slope factor of 0.49 ± 0.04 indicates an evolutionarily conserved coupling ratio of 1:1 H^+^:glutamate exchange. In similar experiments with [Cl^-^]_o_ = 40 mM (Figure S5) we observed smaller shifts of current reversal potential upon changes of the H^+^ Nernst potential that could be fit with a slope factor of 0.29 ± 0.03. Large DVGLUT_PM_ Cl^-^currents that represent a dominant cell conductance reduce voltage shifts due to changes in glutamate transport driving force and thus diminish the slope of the plot of current reversal potential versus H^+^ Nernst potentials.

**Fig. 7.**
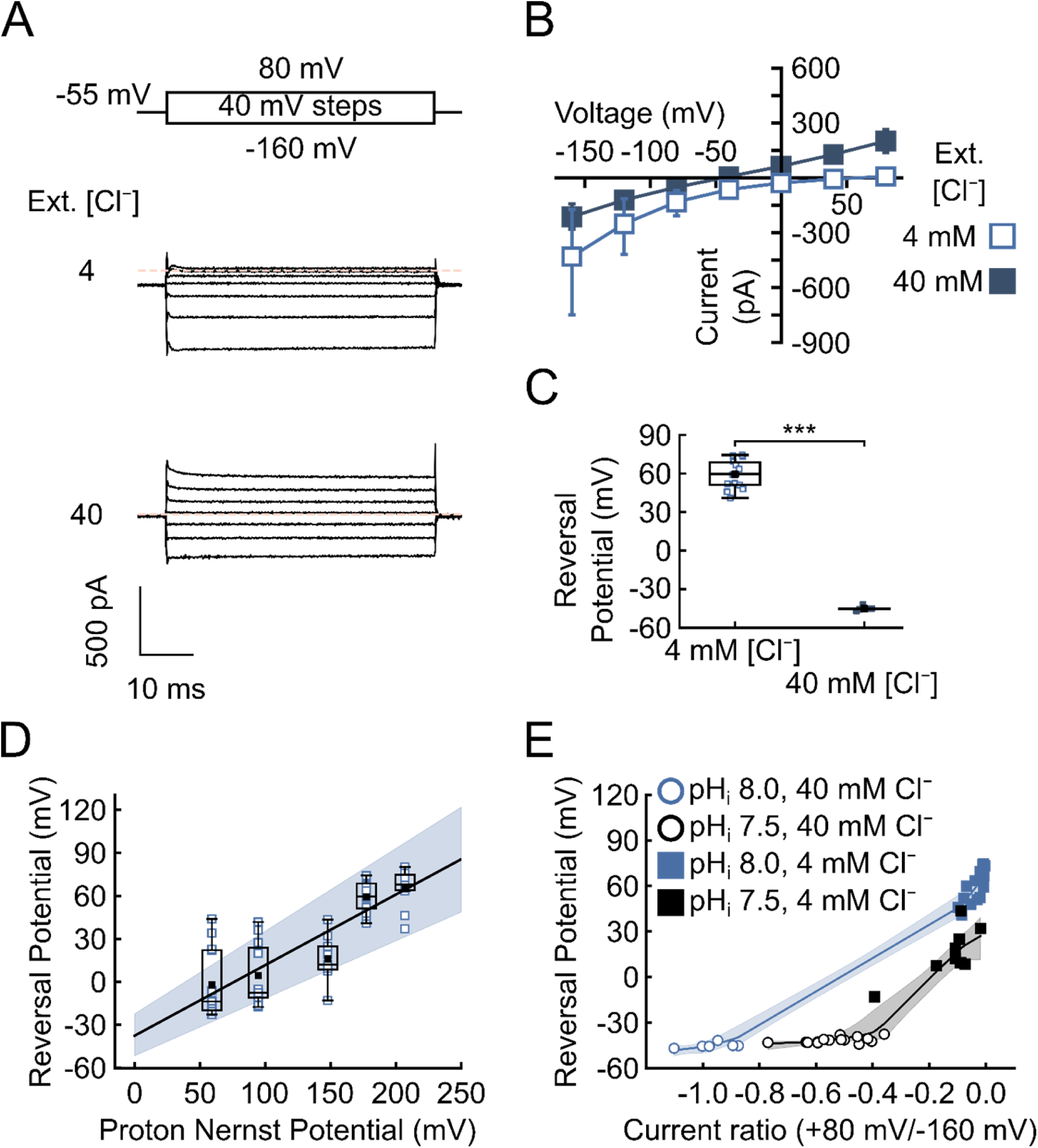
Transport stoichiometry of DVGLUT glutamate transport. **(A)** Representative recordings from two different DVGLUT_PM_-expressing cells dialyzed with 140 mM glutamate at pH_i_ 8.0 each and perfused with the indicated extracellular [Cl^-^] at pH_o_ 5.0. Leak componenets were corrected by subtracting the current values at the same [Cl^-^] at extracellular pH 7.5. Dashed lines indicate zero-current level. **(B)** Current-voltage relationships from experiments as shown in (A). Data are given as means ± 95% CI ([Cl^-^]_o_ **=** 4 mM, n = 14, [Cl^-^]_o_ **=** 40mM, n = 6) **(C)** Current reversal potentials for the two external [Cl^-^]. **(D)** Glutamate reversal potential as a function of Nernst potential for protons H^+^ calculated as 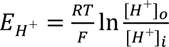. Intracellular pHs = 6.0, 6.6, 7.5, 8.0. 8.5 (n = 13, 11, 9, 14, 10). **(E)** Reversal potentials as function of relative Cl^-^ currents calculated as ratios of current amplitudes elicited at +80 mV and -160 mV for cells dialyzed with internal 140 mM glutamate at pH 8.0 or 7.5. Measurements were performed at 40 mM (open symbols) or 4 mM (closed symbols) [Cl^-^]_o_, both solutions at pH 5.0. Solid lines and shaded areas represent the means and bootstrapped 95% confidence intervals for those fits. Non-overlapping confidence intervals indicate a significant difference between pH_i_ = 8.0 and pHi = 7.5 curves.

As additional test for possible influences of Cl^-^ currents on experimental E_rev_ values, we compared data from experiments with [Cl^-^]_o_ of 4 or 40 mM^12^ that differ prominently in Cl^-^ current amplitudes (Figure 7B) and reversal potentials (Figure 7C). Figure 7D depicts a plot of current reversal potential versus relative Cl^-^ current amplitudes (i.e. ratios of current amplitudes at +80 mV by -160 mV), demonstrating significantly different dependences for pH_i_ = 8.0 and pH_i_ = 7.5. Reversal potentials become more negative with increased relative Cl^-^ current amplitudes at fixed pH_i,_ reaching a pH-independent saturation of reversal potentials at very negative current ratios, which are obviously dominated by VGLUT anion channel currents. However, when comparing two cells with similar relative anion current amplitudes outside this range, reversal potentials at pH _i_ 7.5 were always more negative than at pH_i_ 8.0. We conclude that changes in current reversal potentials report on alterations in the driving force for glutamate transport and that DVGLUT_PM_ functions as coupled H^+^:glutamate exchanger at 1:1 stoichiometry.

### Allosteric Cl^-^ affinity affects glutamate transport rates during vesicle filling

*Drosophila* vesicular glutamate transporters differ from their mammalian counterpart rVGLUT1 in higher allosteric Cl^-^ affinity and larger Cl^-^ currents. To predict how such changes might modify synaptic vesicle filling with glutamate, we used a continuum model (Figure 8), which we recently developed to describe synaptic vesicle filling with a set of differential equations^12^.

**Fig. 8.**
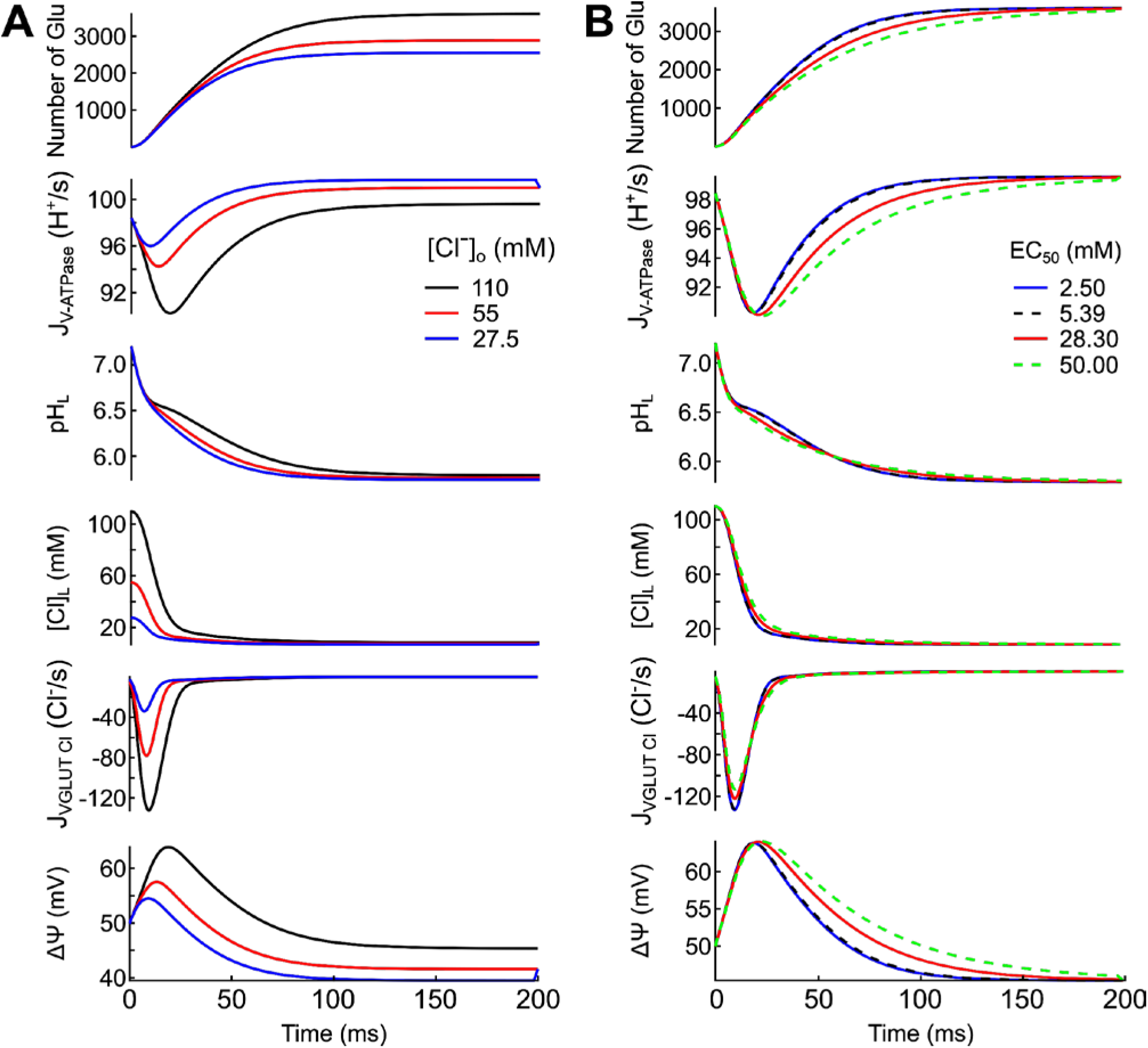
Alteration of vesicular glutamate accumulation in a model glutamatergic synaptic vesicle upon changes in external [Cl^-^] and allosteric affinity. **(A, B)** Predicted evolution of luminal amino acid (AA) numbers, active proton transport rates, pH, [Cl^−^], Cl^-^ currents by VGLUT chloride channels and luminal membrane potential for vesicles formed by endocytosis in external solutions containing 27.5, 55 or 110 mM chloride **(A)**, or with transporters exhibiting three different allosteric affinities at a fixed [Cl^-^] of 110 mM **(B)**.

*Drosophila* haemolymph contains less than 50% of the extracellular [Cl^-^] in mammals^32,33^, so that – directly after endocytosis - DVGLUT will be exposed to much lower luminal chloride concentrations that rVGLUT1. We used the same parameters described for rVGLUT1 ^12^, exclusively modifying the initial [Cl^-^] in the synaptic vesicle. Figure 8A depicts time-dependent changes in luminal glutamate numbers, active proton transport rates, vesicular pH, vesicular [Cl^-^], Cl^-^ currents by VGLUT chloride channels and luminal membrane potential after starting the V-ATPase at time = 0. These calculations were performed for three starting chloride concentrations, 110 mM, 55 mM and 27.5 mM. Reduced starting vesicular [Cl^-^] decreases Cl^-^ outward currents and thus restricts depolarization of the synaptic vesicle and the driving force for glutamate uptake. Lower [Cl^-^] increases ATP consumption and slightly accelerates acidification, with more acidic steady-state pH. Steady-state vesicular [Cl^-^] are only slightly affected by starting values. Figure 8B illustrates how the increased allosteric affinity of DVGLUT partially compensates for this transport impairment. Higher allosteric affinity accelerates glutamate accumulation without affecting steady-state glutamate loading. These changes are associated with higher V-ATPase H^+^ pumping and shorter transient depolarization of the synaptic vesicle.

## DISCUSSION

Here, we studied the *Drosophila* vesicular glutamate transporter using whole-cell patch clamp in transfected mammalian cells and compared its transport functions with rVGLUT1^12^. As expected, rat and fly transporters share many functional features. In the absence of glutamate, both proteins function as anion channels that are activated by extracellular (corresponding to the synaptic lumen) acidification (Figure 1) and extracellular Cl^-^ ions (Figure 3)^11–13^. For *Drosophila* and rat VGLUTs, glutamate transport as well as anion channel opening are stimulated by membrane hyperpolarization (Figures 1 and 2), which corresponds to luminal positive voltages generated by V-ATPase H^+^ pumping in synaptic vesicles. *Drosophila* and rat VGLUT anion channels both exhibit a lyotropic selectivity sequence, with higher current amplitudes for large (I^-^) or polyatomic anions (NO_3_ ^-^) than for Cl^-^ (Figure 4). There are, however, quantitative differences between the two tested transporters. DVGLUT_PM_ activates at lower [Cl^-^] concentrations, with only a few millimolar Cl^-^ being sufficient to fully activate DVGLUT_PM_ anion currents. While single channel amplitudes of 39 ± 5 fA in Cl⁻_i_ were obtained for rat VGLUT1_PM_, we observed much smaller current amplitudes for DVGLUT_PM_ (Figure 5) without difference between Cl^-^ (11 ± 3 fA), NO_3_ ^-^ (11 ± 4 fA) and I^-^ (8 ± 3 fA). Despite the lower unitary current amplitudes, macroscopic DVGLUT anion currents are much larger at comparable expression levels (Figure 6), indicating higher absolute open probabilities of *Drosphila* anion channels than for the rat counterpart. When measured in the absence of glutamate, depolarizing voltage steps after a negative prepulse cause channel deactivation for both transporter-associated anion channels. Anion channel deactivation is complete within few milliseconds in case of rVGLUT1, but it is slower and leaves significant steady-state currents in DVGLUT_PM_ (Figure 2).

We measured glutamate uptake currents in experiments with glutamate as main intracellular anion (Figure 6). To compare unitary transport rates, glutamate uptake currents were measured together with whole-cell fluorescence intensity in each experiment^12,27,28,34^ (Figure 6B). Both VGLUTs were expressed as eGFP-fusion proteins, and whole-cell fluorescences thus report on the number of expressed transporters, in intracellular as well as in plasma membranes. Since DVGLUT_PM_ and rVGLUT1_PM_ exhibit comparable subcellular distributions in confocal images, (Figure S4), and since there were no indications for the saturation of transporter expression and intracellular trafficking (Figure 6B), the numbers of transporters in the surface membrane is expected to be proportional to the whole-cell fluorescence^27,28,34^ . A plot of glutamate currents versus whole-cell fluorescences reveals overlapping distributions for rat and fly transporters (Figure 6B), indicating similar DVGLUT_PM_ transport currents at comparable transporter numbers as rVGLUT1_PM_.

In experiments with glutamate as main intracellular anion, we observed more negative current reversal potentials in cells expressing DVGLUT_PM_ than for rVGLUT1_PM_ (Figure 6D). Such difference in reversal potential may indicate differences in the transport stoichiometries for fly and rat transporters. We recently demonstrated that - under glutamate transport conditions - rVGLUT1_PM_ reversal potentials change linearly with the Nernst potential for H^+^, as expected for a H^+^-glutamate exchanger at 1:1 stoichiometry^12^. When tested at 40 mM external Cl^-^, DVGLUT_PM_ current reversal potentials changed less upon variation of the intracellular pH than values obtained with rVGLUT1_PM_ (Figure S5). However, lowering the external [Cl^-^] to 4 mM, increased the slope of the linear relationship between current reversal potentials and calculated Nernst potentials for H^+^ (Figure 7) to 0.5, indicating a conserved 1:1 H^+^:glutamate transport stoichiometry also for DVGLUT_PM_ ^12^. We conclude that fly and rat vesicular glutamate transporters resemble each other in glutamate transport rates and transport stoichiometry.

In cells internally perfused with glutamate, currents are entirely carried by glutamate transport at voltages negative to the reversal potential, whereas Cl^-^ currents conducted by VGLUT anion channels can be observed at positive voltages (Figure 6). The comparison of Cl^-^ currents at positive potentials in experiments with (Figure 6) or without intracellular glutamate (Figure 1) reveals an important functional effect of glutamate on the VGLUT anion channel function. Intracellular glutamate keeps the anion channel open to permit Cl^-^ influx from the extracellular space in our experiments and from the synaptic vesicle lumen under physiological conditions^12^.

When measured under conditions permitting glutamate uptake, DVGLUT_PM_ Cl^-^ currents were much larger than rVGLUT1_PM_ Cl^-^ outward currents at comparable whole-cell fluorescences (Figure 6). Since DVGLUT_PM_ unitary currents are smaller than those of the rat transporters (Figure 5C), anion channel open probabilities need to be significantly larger for the fly than for the rat transporter. The higher anion channel open probability is a major functional difference between vesicular glutamate transport of the two species.

Martineau et al.^35^ proposed that VGLUT chloride channels mediate Cl^-^ efflux from synaptic vesicles to ensure osmotic neutrality of synaptic vesicle filling. Subsequent mathematical modeling demonstrated that changes in the VGLUT Cl^-^ conductance modify the speed and the final levels of glutamate filling^12^. We currently imagine the following order of transport events during synaptic vesicle filling^12,36^: immediately after endocytosis synaptic vesicles contain high Cl^-^ levels of the extracellular solution. V-ATPase activity acidifies synaptic vesicles and activates both VGLUT transport functions. Since VGLUT anion channels exhibit higher Cl^-^ transport rates than VGLUT glutamate transport, the first consequence of synaptic vesicle acidification is the rapid depolarization of the synaptic lumen by Cl^-^ outward permeation through the VGLUT channel, which increases the driving force of glutamate accumulation. Concurrent glutamate accumulation and Cl^-^ efflux exchange Cl^-^ to glutamate as main anion in synaptic vesicles. Because of the large driving force of H^+^-glutamate exchange, glutamate accumulation does not reach equilibrium conditions, but might rather be terminated by allosteric deactivation at low vesicular [Cl^-^].

This scenario links the differences of rat and fly VGLUT properties in the affinity of the allosteric Cl^-^ binding site and in anion channel activity to variation in electrolyte concentrations in the extracellular space of fly and rat neurons. The *Drosophila* haemolymph contains less extracellular [Cl^-^] (between 30 mM^32^ and 42.2 mM^33^) than mammalian extracellular solutions, so that – directly after endocytosis – DVGLUT will be exposed to much lower luminal chloride concentrations that rVGLUT1. We used a continuum model for synaptic vesicle filling^12^ to illustrate possible effects of lower external [Cl^-^] and separate allosteric Cl^-^ activation on synaptic vesicle glutamate accumulation. Without altering unitary currents, absolute open probabilities and numbers of VGLUT anion channels, lower external [Cl^-^] reduces Cl^-^ currents, restricts depolarization of the vesicle and reduces the speed and the steady-state glutamate accumulation at slightly increased ATP consumption (Figure 8A). Higher anion channel open probabilities of DVGLUT will depolarize the synaptic vesicle more effectively and enhance the initial glutamate loading rate. However, there are only small effects on overall glutamate loading upon sole variation of VGLUT anion conductances^12^. Increasing the allosteric site affinity accelerates glutamate uptake (Figure 8B), thereby reducing V-ATPase activity and thus ATP consumption. This mechanisms might partially compensate for the effects of the low Cl^-^ concentrations in the *Drosophila* hemolymph on glutamate accumulation.

VGLUTs combine carrier and channel functions^10–12,36^. While the importance of VGLUT glutamate transport for presynaptic glutamate release is evident^37,38^, the cellular roles of VGLUT anion channel functions are not as obvious. The comparison of vesicular glutamate transporters of two species demonstrates major differences in allosteric regulation by anions and in anion channel function, but not in glutamate transport, which is the main physiological function of these transporters. The adaptation of these two functional VGLUT features to differences in ionic conditions underlines the impact of the VGLUT Cl^-^ channel function and the allosteric activation by Cl^-^ for normal synaptic vesicle filling.

## EXPERIMENTAL MODEL AND STUDY PARTICIPANT DETAILS

### Generation of PAC KO cells

Knock-down of human PACC1 (gene encoding for PAC/TMEM206^22,23^ in the embryonic kidney cell line HEK293T was performed with Crispr-Cas9 technique^39^. We designed target-specific guidance RNAs with a web-based design tools. (https://www.benchling.com/crispr/) : (1a) Top Strand Target (Exon 3 Target): 5’ACGGTCAGAGTCCAAGGTCC 3’ and (1b) Bottom Strand Target (Exon 3 Target): 5’GCTCTGAGTTCTCAACCACC 3’ (2a) Top Strand Target (Exon 4 Target): 5’CTACCTGCTGCTCATGGCTG 3’ and (2b) Bottom Strand Target (Exon 4 Target): 5’GCAGGTAGATGAAGATGAGT 3’ Cas9 nickase (Cas9n) was used to introduce nicks on both strands of the target DNA in Exon 3 and 4. Plasmids used for Cas guidance (pSpCas9n(BB)-2A-GFP (PX461):#48140, pSpCas9n(BB)-2A-Puro (PX462) V2.0.:#62987) were obtained from Addgene (www.addgene.org). The resulting double strand breaks are reconnected by cellular nonhomologous end joining. Single clones were isolated after antibiotic selection with puromycin and FACS sorting for GFP-positive single cells. Knock-out of PACC1 was verified by PCR amplification and subsequently sequencing of Exon 3 and 4 in PACC1 (Eurofins Genomics Germany GmbH, Ebersberg) with the following primers. Primers: O.38965’GGTTTTTGGTGACAATTAAGTGACAAGCAGC 3’ S TMEM206 Exon3 gDNA SeqFor O.38975’CAGCCTAGACATATCCTCTGTTCCAAG 3’ AS TMEM206 Exon3 gDNA SeqRev O.38985’CCTCCTTGGTGAGGCAGGCAAG 3’ S TMEM206 Exon4 gDNA SeqFor O.38995’GTAGTACCACAAATACAACGGCAGCTC 3’ AS TMEM206 Exon4 gDNA SeqRev Clones identified by PCR were further tested by electrophysiological assays. KO_PAC_HEK293T cells show whole-cell resistances of 4.2 ± 0.2 GΩ over a voltage range between -200 mV and 150 mV at pH_o_ 7.4 and 4.4 ± 0.3 GΩ at pH_o_ 5.5 (Figure S2).

## METHOD DETAILS

### Construction of expression plasmids, mutagenesis and heterologous expression

The nucleotide sequence harboring the gene encoding the *Drosophila* VGLUT transporter was obtained from Genbank (accession no. NM_001258919.1). First strand cDNA synthesized on *Drosophila* head mRNA was used to amplify the coding sequence of DVGLUT. The forward primer (5’-AGCTGAAGCTTCCACC**<u>ATG</u>**AAGGGTCTGACGGCG-3’) introduced a *Hind*III restriction site at the 5’ end followed by a Kozak (1984; CCACC) consensus motif preceding the initiating ATG codon. The reverse primer (5’-CGTTATCTAGACTGCGTCTGGTATCCCTG-3‘) introduced a unique *Xba*I restriction site at the 3’ end. Amplified fragments were purified using a Nucleo Spin kit (Macherey-Nagel; Thermo Fisher Scientific, Dreieich, Germany). Fragments were digested with *Hind*III and *Xba*I. After agarose gel electrophoresis, fragments of expected length were purified (Nucleo Spin kit) and ligated into *Hind*III/*Xba*I cut pBluescript (SK-) vector (Stratagene; Agilent Technologies, Waldbronn, Germany) for sequencing (Eurofins Genomics Germany GmbH, Ebersberg). The sequence verified cDNA was subsequently transferred into *Hind*III/*Xba*I cut pcDNA3.1(+) vector (Invitrogen; Thermo Fisher Scientific, Dreieich, Germany) for expression in eukaryotic cells. To express DVGLUT as an eGFP fusion protein, we inserted the coding region of eGFP into pcDNA3.1-DVGLUT at the 3ʹ end of the transporter coding region.

To promote surface insertion, residues Y38, E39, E40, M41, E42, G43, and G44, which form an acidic stretch near the N terminus, were substituted for alanine using PCR-based mutagenesis. Transient expression of the resulting mutant (referred to as DVGLUT_PM_) led to predominant localization at the plasma membrane (Figure S1). DVGLUT_PM_ was studied in three expression systems: KO_PAC_HEK 293T cells (see above) and HEK 293T cells (Sigma-Aldrich) that were transiently transfected with 2–5 μg plasmid DNA using a lipofectamine-based method and examined 36 h post-transfection, and stably transfected Flp-In T-REx 293 cells that were induced with at least 1.5 μg ml⁻¹ tetracycline for up to 24 hours ^40^.

### Whole-cell patch clamp and fluorescence-current correlation

Standard whole-cell patch clamp recordings were performed using an EPC-10 amplifier, controlled by PatchMaster (HEKA, Germany)^13^. Borosilicate pipettes (Harvard Apparatus, USA) with resistances < 2.5 MΩ were used for experiments with pipette solutions containing high concentrations of small intracellular anions (chloride, nitrate, iodide), and < 3.5 MΩ with high glutamate concentrations. In all cases, series resistance compensation limited voltage errors to < 5 mV.

The standard bath solution for channel current recordings contained (in mM): 136 choline chloride, 1 CaCl_2_, 1 MgCl₂, 10 HEPES or MES, with osmolarity adjusted with glucose to be 10–15 mOsm higher than that of the pipette solution. The extracellular pH was adjusted with choline hydroxide (CholOH) to values between 6.5 and 8.2 using HEPES as buffer and for more acidic values using MES as buffer. Titration with extracellular chloride (Figure 3) was performed by stepwise substitution of chloride with gluconate at pHₒ 5.0 with 50 mM MES. For experiments quantifying transport currents at 4 mM Cl^-^ choline, Cl^-^ was replaced with a mixture of 136 mM choline hydroxide and glutamic acid; for experiments with 40 mM Cl^-^ 36 mM choline-glutamate with substituted with 36 mM choline-chloride. In experiments with lower external [Cl^-^] the holding potential was set negative to the reversal potential, such that baseline currents were below the zero-current level.

The standard pipette solution contained (in mM) 125 choline chloride, 2 MgCl₂, 5 EGTA, 10 HEPES, pHᵢ 7.4. To study unitary current amplitudes for different intracellular anion compositions (Figure 5C), choline chloride was equimolarly substituted with choline iodide or cholineNO_3_, or partially substituted with choline glutamate; all solutions were titrated using tetramethylammonium hydroxide (TMAOH) or choline hydroxide (CholOH) to pH_i_ 7.4. For experiments, in which fluorescence–current relationships were generated with cells dialyzed with internal I^-^- or NO ^-^-based solutions, the concentration of the sole charge-carrying anion was 140 mM with 5 mM MgCl₂ replaced by 5 mM Mg(OH)₂. For estimation of coupling stoichiometry, at pHᵢ values below 6.5, 30 mM HEPES was replaced with 30 mM MES and at higher pHᵢ values with 30 mM AMPSO; all solutions were titrated using CholOH. After establishing the whole-cell mode recordings were started within the first two minutes.

In experiments combining fluorescence measurements and whole-cell recordings (Figures 1C and 6B), cells were imaged before establishing the whole-cell mode to prevent modification of the subcellular distribution of VGLUTs by intracellular anions. eGFP was excited using a Polychrom V monochromator (TILL Photonics) set to 488 nm and recorded using a Neo camera (Andor)^27^. Mean fluorescence intensities for manually selected regions of interest were analyzed using Fiji (http://www.fiji.sc).

### Noise analysis

Currents were digitized at 100 kHz during 50s voltage steps to –100 mV at pHₒ 7.4 and subsequently at pHₒ 5.5, using 45 s fragments for analysis. Power spectral densities (PSDs) were computed from mean-detrended traces using Welch’s method (Hann window, ∼2 Hz frequency resolution, 50% overlap). For each cell, the PSD obtained under neutral conditions was subtracted from that recorded at acidic pH to generate a difference spectrum. The resulting spectra were limited to 2–8 kHz and log-binned (60 bins per decade). At each frequency bin, spectra were averaged across all cells. The aggregated spectrum was then fitted with pink noise (1/fᵃ) plus one or two Lorentzian components, and Akaike information criterion (ΔAIC > 6) was used to determine whether a double-Lorentzian provided a significantly better fit:

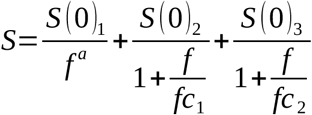

with S denoting the spectral density, f the frequency, a the slope of the linear pink noise, and S(0) and fc the amplitude and the corner frequencies respectively.

Unitary current amplitudes were determined from current fluctuations measured between 99 and 149 ms after the voltage step at –160 mV, using 5–8 extracellular solutions with pH values between 5.0 and 8.2. Mean currents were taken from unfiltered traces and high-frequency noise was isolated by high-pass filtering to approximately 100 Hz to remove slow relaxations in the current traces. The resulting variance-mean current plots did not cover a full parabola, so F-tests (p < 0.01) were used to assess whether adding a quadratic term significantly improved the fit compared with a linear model for each dataset acquired under a given charge carrier, defined by the cytosolic anion used for dialysis. When the quadratic term improved the fit, unitary current amplitudes were calculated from parabolic fits. This was only necessary for rVGLUT1_PM_ data, for DVGLUT, linear fits were sufficient. A global fit with one shared single-channel amplitude and an independent **N** term for each cell was applied to all datasets under the same condition. When the F-test did not support the quadratic model, unitary current amplitudes were instead derived from global linear fits of the form:

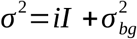

Current-variance relationships (Figure 5) were obtained from resampled variance–mean data using bootstrap resampling with replacement, and distributions of unitary current amplitudes were generated for each condition.

pH dependences (Figures 1 and 4) were obtained by varying the extracellular pH during whole-cell recordings and plotting normalized steady-state currents measured at −160 mV as a function of pH. Current responses did not saturate within the tested pH range; therefore, a phenomenological hyperbolic-type model was used to describe the pH dependence:

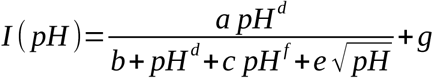

Bootstrap resampling of current responses was used to generate distributions of fitted curves, and pH dependences were considered significantly different when the corresponding bootstrap distribu - tions did not overlap, except at extreme pH values.

### Confocal imaging

Confocal imaging of living KO_PAC_HEK293T cells bathed in CholineCl was performed using an inverted confocal laser scanning microscope (Leica Stellaris 5, Leica Microsystems, Heidelberg, Germany) equipped with a white light laser. Samples were excited at 488 nm and emission was collected between 490–600 nm. Laser power and detector gain were optimized to 2% and 15% respectively and kept constant across all images for both DVGLUT_PM_ and rVGLUT1_PM_ constructs for quantitative comparison. Images were acquired using a 63×/1.4 oil-immersion objective at a resolution of 2048 × 2048 pixels scanned at 600Hz. Eight-bit images were collected with a frame averaging of 4. Image analysis was performed using FIJI (ImageJ) (Figures S4C and S4D): for each cell, intensity values were measured at ten pixel along the plasma membrane and compared to five intracellular area-averaged intensity measurements. Figure S4C provides a plot of average membrane values versus the sum of membrane and cytoplasmic fluorescence intensities for each analyzed cell. Ratios of membrane by total fluorescences were used to quantify relative plasma membrane insertion of fly and rat VGLUTs in Figure S4D.

### Modeling glutamate accumulation in synaptic vesicles

We model the consequences of changes in external [Cl^-^] and allosteric affinity using a recently developed continuum quantitative model of synaptic vesicles^12^ that describes the temporal evolution of luminal glutamate/aspartate molecule numbers, pH, and [Cl^-^] with a set of differential equations. The model contains V-type ATPases as the only source of acidification. The two VGLUT transport modes are represented by two equations, one for H^+^-glutamate exchange and an additional one for the chloride channel; both change with luminal pH with the same pK_M_ (5.5) and on luminal [Cl^-^] with the same concentration dependence. Open probabilities of VGLUT Cl^-^ channels and maximum transport rates for VGLUT glutamate transporters were assumed to be voltage-independent. Proton leak channels are required to describe constant steady-state vesicular pH during sustained V-type ATPase proton pumping.

Mathematical modeling starts with ion concentrations of the external solution: neutral pH and negligible glutamate concentrations. We varied the external [Cl-] between 27.5 mM and 110 mM. Intracellular cytoplasmic [Cl^-^] was set to 21 mM^41^ and cytoplasmic [glutamate] and [aspartate] to 10 mM^42^.

### Statistical analysis

Unless noted otherwise, statistical comparisons were performed using bootstrap-based non-parametric tests, with * denoting p < 0.05, ** < 0.01, and *** < 0.001. In the plots, error bars represent the non-parametric bootstrap percentile 95% confidence interval of the mean. In the text, values are reported as mean ± SEM.

F-tests were used to assess whether adding parameters improved the quality of model fits. Comparisons were performed between flat (zero-slope) and sloped linear fits for current–expression relationships, and to assess whether parabolic models provided a significantly better description of the variance–mean current relationships than linear ones. A significance threshold of p < 0.01 was applied, with higher values considered not significant.

## ACKNOWLEDGMENTS

We thank Dr. Achmed Mrestani and Manfred Heckmann for helpful discussions. This work was supported by the Deutsche Forschungsgemeinschaft (German Research Foundation) to Ch.F. (FA 301/15–2) as part of Research Unit FOR 2518, *DynIon*; to Ch.F. (FA 301/13-1 and 15–2) and G.U. as part of the Research Unit FOR 2795.

## DECLARATION OF GENERATIVE AI IN SCIENTIFIC WRITING

During the preparation of this manuscript, the authors used Grammarly to assist with grammar, spelling, and syntax. This writting assistance tool was used with the sole purpose of improving readability and it was applied under full human oversight. All content was reviewed, edited, and verified by the authors, who take full responsibility for the content of the published article.

## AUTHOR CONTRIBUTION

Conceptualization, V.L. and Ch.F.; data curation, V.L., Y.G., S.U.,;funding acquisition, G.U., Ch.F.; investigation, V.L., Y.G., S.U., G.U.; Ch.F.; methodology, A.F., S.B., A.B..; writing – origi - nal draft, V.L., A.B. and Ch.F.; writing – review & editing, V.L., G.U., Ch.F.

