## Supplemental Figures for "Evolutionary optimization of allosteric activation by Cl^-^ and Cl^-^ conduction in vesicular glutamate transporters"

# A

**Amino terminus**

```
DVGLUT       MKGLTAFKEKATGVFGGLKPNMEKFEISQSYHGGHGGYEEMEGGDREGRGPGGGHAYDDD 60
hVGLUT1      ------------------------MEF-----------RQEEFRKLAGRALGKLHRLLEK 25
rVGLUT1      ------------------------MEF-----------RQEEFRKLAGRALGRLHRLLEK 25
                                     :*:           .: *  .  **. *  *   :.
```

# B

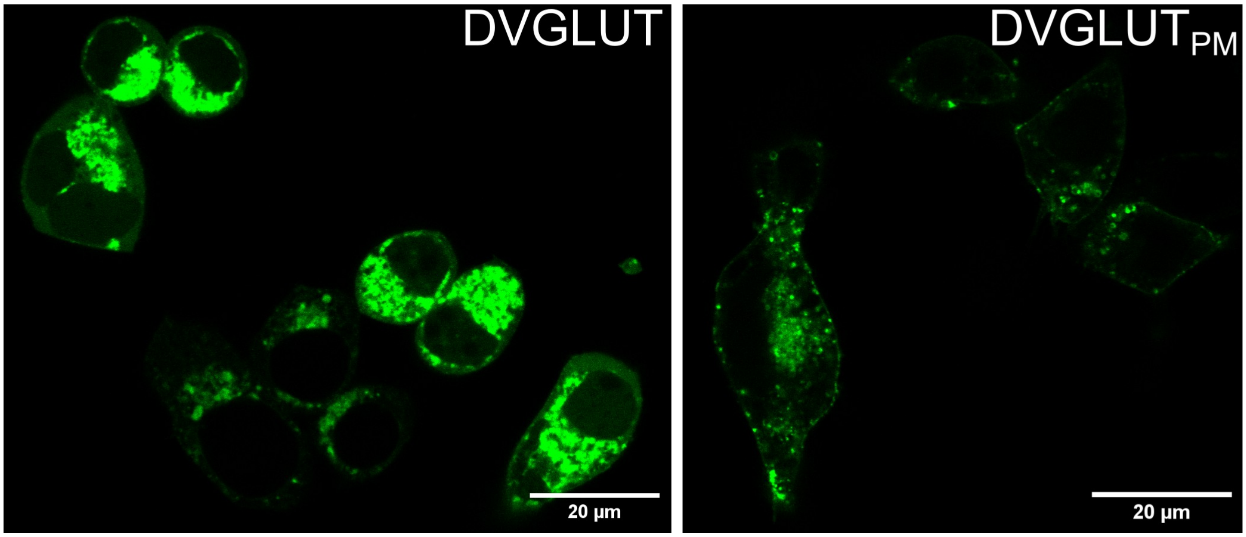


**Fig. S1. Mutations in the amino-terminus improve surface membrane insertion of DVGLUT.**

**(A)** Sequence alignment of the amino termini of *Drosophila* vesicular glutamate transporter (DVGLUT), human VGLUT1 (hVGLUT1), and rat VGLUT1 (rVGLUT1). Alanine substitutions were introduced at the highlighted residues into the DVGLUT sequence. **(B)** Confocal images of $KO_{PAC}$HEK293T cells transiently expressing DVGLUT (left) or plasma membrane optimized $DVLGUT_{PM}$ (right).

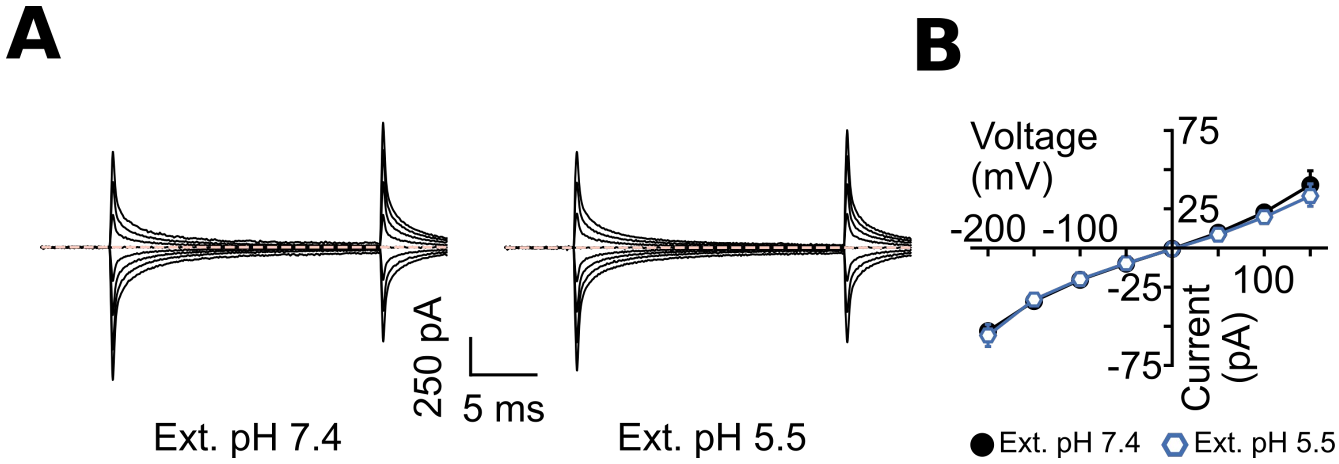


**Fig. S2. Background conductances in $KO_{PAC}$HEK293T cells.**

**(A)** Representative whole-cell chloride current recordings from $KO_{PAC}$HEK293T cells perfused with $Cl^-$-based solutions at $pH_o$ 7.4 or $pH_o$ 5.5 in response to voltage steps from −200 mV to +150 mV in 50 mV increments. **(B)** Current-voltage relationships from untransfected $KO_{PAC}$HEK293T cells perfused with $Cl^-$-based solutions at $pH_o$ 7.4 or $pH_o$ 5.5 in response to voltage steps from −200 mV to +150 mV in 50 mV increments (*n* = 10).

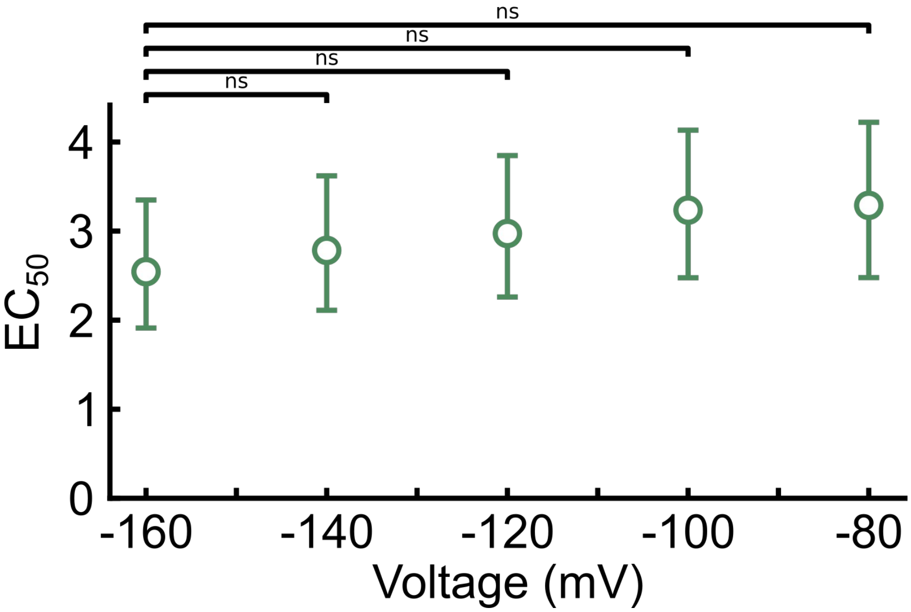


**Fig. S3. $EC_{50}$ for external chloride are voltage-independent.**

For each voltage, dose–response relationships for external chloride were fit using a Hill-type equation with cooperativity fixed to 1: $I([Cl^{-}]_o) = I_{min} + (I_{max} - I_{min})\ [Cl^{-}]_o^{\,n} / EC_{50}^{\,n} + [Cl^{-}]_o^{\,n}$. The resulting $EC_{50}$ values are plotted as a function of membrane potential. Symbols represent mean values, and error bars indicate the 95% CI. Differences across voltages were not significant.

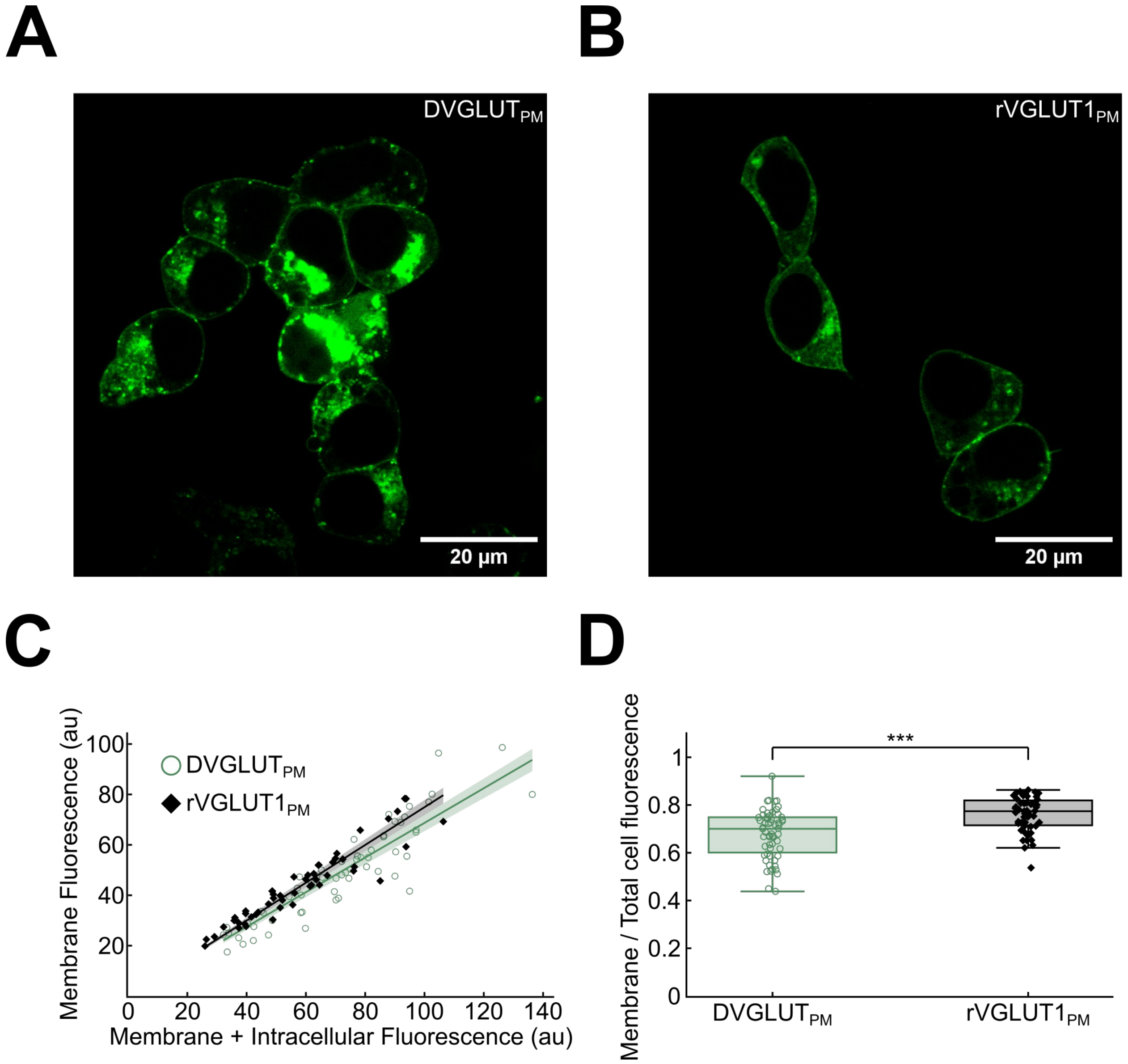


**Fig. S4. Membrane insertion of rat and fly mutant transporters**

**(A, B)** Representative confocal images of $\text{KO}_{\text{PAC}}$HEK293T cells either transiently expressing $\text{DVGLUT}_{\text{PM}}$ (left) or $\text{rVGLUT1}_{\text{PM}}$ (right). (C) Plots of membrane bound fluorescence and total fluorescence from cells expressing $\text{DVGLUT}_{\text{PM}}$ ($n = 59$, open symbols) or $\text{rVGLUT1}_{\text{PM}}$ ($n = 51$, closed symbols). (D) Relative membrane insertion for $\text{DVGLUT}_{\text{PM}}$ (open symbols) or $\text{rVGLUT1}_{\text{PM}}$ (closed symbols).

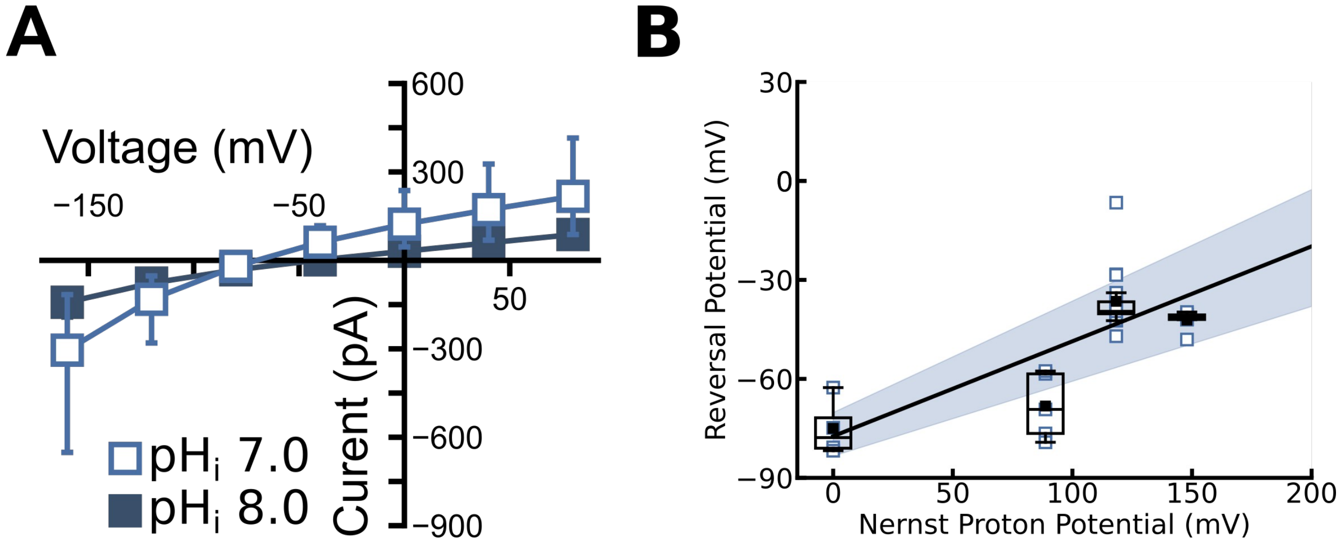


**Fig. S5. The DVGLUT$_{PM}$ anion conductance affects the apparent pH dependence of current reversal potentials in the glutamate transport mode.**

**(A)** Current-voltage relationships from experiments with DVGLUT$_{PM}$-expressing cells dialyzed with 140 mM glutamate at indicated pH$_i$ and perfused with a solution containing [Cl$^-$] = 40 mM. Data are given as means ± 95% CI (pH$_i$ = 7.0, n = 5, pH$_i$ = 8.0, n = 6). **(B)** Changes of current reversal potentials of cells intracellularly dialyzed with glutamate-based solution and perfused with an external solution containing 40 mM Cl$^-$ (n = 4 (pH 5.5), n = 5 (pH 7.0), n = 17 (pH 7.5), n = 6 (pH 8.0) cells) with the Nernst potential for H$^+$ calculated as $E_H = \frac{RT}{F}\ln\frac{[H^+]_o}{[H^+]_i}$. Lines and shaded areas show the mean and 95% CI from linear fits.
